# Propiconazole and Tianfengsu Regulated the Growth, Flavor, and Nutrition *of Brassica rapa* by Brassinosteroid Pathway

**DOI:** 10.1101/2024.05.06.592836

**Authors:** Dekang Guo, Qing Gao, Yunxue Song, Zhicheng Liu, Daorui Wang, Hanhong Xu, Fei Lin

**Affiliations:** National Key Laboratory of Green Pesticides/Key Laboratory of Natural Pesticide and Chemical Biology, Ministry of Education, South China Agricultural University, Guangzhou, 510642, Guangdong, China; Guangzhou Academy of Agricultural and Rural Sciences, Guangzhou 510335, Guangdong, China; Jiangmen Daguangming Agricultural Chemical New Association Co., Ltd. Jiangmen, 529000, Guangdong, China

**Keywords:** *Brassica rapa*, Propiconazole, Tianfengsu, Brassinosteroids, Flavor quality, Weighted gene co-expression network analysis

## Abstract

Propiconazole (PCZ) and Tianfengsu (TFS) are widely used plant growth regulators in vegetable production for improving crop growth, quality, and flavor. This study investigated the effects of PCZ and TFS, applied individually or in combination, on the growth, development, flavor quality, and nutritional components of choy sum (*Brassica rapa*) and *Arabidopsis thaliana*, as well as the underlying molecular mechanisms. The results showed that PCZ inhibited the growth of choy sum and *Arabidopsis* but enhanced the accumulation of flavor compounds such as soluble sugars, proteins, and vitamin C. In contrast, TFS promoted plant growth and increased the content of nutritional components, including chlorophyll and unsaturated fatty acids. Notably, the combined application of PCZ and TFS significantly improved overall plant quality, achieving the optimal balance of flavor and nutritional value while maintaining high yield. Transcriptomic analysis revealed the molecular mechanisms of PCZ and TFS in differentially regulating the expression of brassinosteroid (BR) signaling and downstream metabolism-related genes. Weighted gene co-expression network analysis (WGCNA) further identified key gene modules and hub genes controlling flavor metabolism in choy sum. This study elucidates the synergistic mechanisms of PCZ and TFS in regulating vegetable growth and quality formation, providing valuable insights for the safe production of high-quality choy sum and the development of novel plant growth regulators or elite varieties.

**HIGHLIGHTS:**

- PCZ and TFS treatments differentially modulate choy sum growth and development by regulating the BR pathway, with PCZ inhibiting while TFS promoting growth.
- PCZ enhances the accumulation of soluble sugars, soluble proteins, and vitamin C, while TFS increases photosynthetic pigments and unsaturated fatty acids, synergistically improving the flavor and nutritional quality of choy sum.
- Transcriptomic analysis and WGCNA uncover key genes and modules controlling flavor metabolism in choy sum, providing potential targets for developing novel plant growth regulators or breeding elite varieties

## 1. Introduction

Choy sum (*Brassica rapa*) is a widely consumed leafy vegetable in southern China, favored for its unique flavor and high nutritional value [1,2]. The eating quality of choy sum, especially the flavor, is a key factor affecting consumer preference and market value [3]. The dwarf plant architecture, characterized by its small leaves and thick stems, is favor by the consumer and facilitates efficient bulk harvesting [4]. To meet this demand, propiconazole (PCZ), a triazole fungicide, is frequently applied off-label to inhibit the vegetative growth of choy sum, to obtain the compact stature and dark green color [5,6]. Although PCZ application can effectively control the plant size, its potential effects on the flavor quality of choy sum, as well as its potential dietary risk remain largely unknown.

Previous studies have shown that PCZ can interfere with the biosynthesis of brassinosteroids (BRs), an important class of plant hormones, by specifically inhibiting the cytochrome P450 monooxygenases involved in the BR biosynthetic pathway [7,8]. BRs are a group of polyhydroxylated steroidal phytohormones that play essential roles in regulating various aspects of plant growth and development, including cell elongation, cell division, vascular differentiation, senescence, and stress responses [9,10]. BR deficiency or insensitivity can lead to typical phenotypes such as dwarfism, dark green leaves, delayed flowering and reduced fertility [11]. In addition to their essential roles in regulating plant growth and development, BRs have also been reported to improve the nutritional quality of several vegetable crops. For example, exogenous application of 24-epibrassinolide (EBR), a bioactive BR, was found to increase the contents of glucosinolates and total phenolics in Chinese kale [12]. Similarly, foliar spray of EBR enhanced the accumulation of ascorbic acid and soluble sugars in tomato fruits [13]. These findings suggest that BRs may play important roles in regulating the biosynthesis and accumulation of flavor compounds in vegetables. However, the molecular mechanisms underlying the effects of PCZ-mediated BR inhibition on vegetable flavor quality remain elusive.

Tianfengsu (TFS) is a commercially available formulation consisting of two bioactive brassinosteroids (BRs), namely 22R,23R,24R-28-epiCS (22R,23R,24R-EBL) and 24-epiCS (24-EBL). These BRs have been extensively studied and shown to enhance plant growth and improve stress tolerance in various crops [14,15]. In recent years, TFS has been widely used in combination with PCZ, aiming to mitigate the growth-inhibiting and quality-compromise caused by excessive application of PCZ [16]. However, it remains unclear the mechanism underlying the interaction between TFS and PCZ during regulating the plant architecture and flavor quality.

In the present study, we aimed to investigate the effects of PCZ and TFS treatments on the growth and flavor quality of choy sum, and elucidate the underlying molecular mechanisms by integrating transcriptomic and metabolomic analyses in *Arabidopsis thaliana*. We treated the choy sum plants with PCZ and TFS at the heading stage, and evaluated their effects on plant growth parameters and the contents of major flavor compounds, including soluble sugars, organic acids, amino acids and fatty acid. Furthermore, we performed comparative transcriptomic and metabolomic profiling on the control and PCZ-treated choy sum leaf samples, and constructed a weighted gene co-expression network to identify the key genes and metabolites involved in flavor biosynthesis pathways. Our findings provide novel insights into the molecular mechanisms of PCZ-mediated flavor modification in leafy vegetables, and shed light on the role of BRs in regulating vegetable flavor quality. The results of this study may facilitate the development of new strategies for improving the flavor quality of choy sum and other leafy vegetables, such as marker-assisted breeding and rational application of plant growth regulators.

## 2. Materials and Methods

### 2.1. Plant materials and growth conditions

Choy sum (*B. rapa* var. parachinensis cv. Sijiu-19) and *Arabidopsis thaliana* (ecotype Col-0) were used in this study. Choy sum seeds were obtained from the Guangzhou Vegetable Research Institute, and *Arabidopsis* seeds were purchased from the *Arabidopsis* Biological Resource Center (ABRC, https://abrc.osu.edu/). Seeds were surface-sterilized with 75% ethanol, stratified at 4 ℃ for 2 days, and sown in a steam-sterilized (121 ℃, 5 h) mixture of vermiculite, peat moss, and perlite (1:3:1, v/v/v). Seedlings were grown in a controlled environment with a 14-h light/10-h dark photoperiod, day/night temperatures of 22 ℃/20 ℃, a light intensity of 300 μmol·m-2·s-1, and 70% relative humidity. Two-week-old seedlings were transplanted into pots containing fresh substrate and continued to grow under the same conditions.

### 2.2. Preparation and application of chemical solution

PCZ (97% purity) and TFS (a mixture of 0.0034% 24-epibrassinolide and 0.0066% 22,23,24-epibrassinolide, 0.01% active ingredients) were provided by the National Key Laboratory of Green Pesticides (Guangzhou, China) and Jiangmen Agrochemical Co., Ltd. (Jiangmen, China), respectively.

PCZ and TFS stock solutions were prepared in Dimethylsulfoxide (DMSO) and diluted with water to working concentrations of 0.15 mM and 0.01 μM, respectively. The combined PCZ+TFS treatment was prepared by mixing the two working solutions at a volume ratio of 1:3 (TFS:PCZ, v/v). A mock solution containing an equal volume of DMSO (0.1%, v/v) was used as the mock control.

Four-week-old choy sum or *Arabidopsis* with pre-bolting was randomly assigned to one of four groups: PCZ (0.15 mM), TFS (0.01 μM), PCZ+TFS and a mock control. Each group comprised 11 plants, and the experiment was replicated three times using a randomized design. *Arabidopsis* plant was irrigated with 20 mL of the corresponding solution, while each choy sum plant was sprayed with 10 mL. After 7 days of treatment (DAT), two types of leaves were collected from choy sum plants for morphological and physiological analyses: the largest rosette leaves, which exhibited deformed morphology likely due to the chemical treatments (referred to as “deformed leaves”, DL), and the newly developed leaves subtending the flower buds (referred to as “Qi Kou leaves”, QKL). Choy sum plants were then subjected to plant architecture observation and flavor-related compounds measurement, while *Arabidopsis* plants underwent plant architecture observation and transcriptomic analyses.

### 2.3. Determination of soluble sugar content

Soluble sugar content was determined using the anthrone-sulfuric acid method with slight modifications [17]. Briefly, 0.5 g of liquid nitrogen-ground sample was extracted with 10 mL of distilled water in a boiling water bath for 1 h. The extract was cooled and adjusted to 25 mL. A 0.1-mL aliquot was mixed with 0.5 mL of anthrone-ethyl acetate reagent and 5 mL of concentrated sulfuric acid. After incubation at room temperature for 10 min, the absorbance at 630 nm was measured using a SHIMADZU UV-1780 spectrophotometer (Shimadzu Corporation, Kyoto, Japan). Soluble sugar content was calculated based on a glucose standard curve (0-100 μg·mL^-1^).

### 2.4. Determination of soluble protein content

Soluble protein content was measured using the Coomassie Brilliant Blue G-250 staining method [18]. A 0.5-g sample was homogenized in 5 mL of extraction buffer (0.1 M Tris-HCl, pH 8.0) and centrifuged at 12,000 ×g for 15 min at 4 ℃. The supernatant (0.1 mL) was mixed with 5 mL of Coomassie Brilliant Blue G-250 staining solution and incubated for 5 min. The absorbance at 595 nm was measured, and soluble protein content was calculated using a bovine serum albumin (BSA) standard curve (0-100 μg·mL-1).

### 2.5. Determination of reduced ascorbic acid content

Reduced ascorbic acid (AsA) content was determined using the molybdenum blue method [19]. One gram of fresh sample was ground in 10 mL of 1.0% oxalic acid-EDTA solution and adjusted to 50 mL. A 10-mL aliquot was sequentially mixed with 1 mL of 2% phosphomolybdic acid, 2 mL of 5% sulfuric acid, and 4 mL of ammonium molybdate solution. After standing for 15 min, the absorbance at 705 nm was measured, and AsA content was calculated based on an AsA standard curve (0-100 μg·mL^-1^).

### 2.6. Determination of chlorophyll and carotenoid contents

Chlorophyll a (Chl a), chlorophyll b (Chl b), and carotenoid (Car) contents were determined using the acetone-ethanol method [20]. A 0.5-g leaf sample was extracted with 20 mL of acetone-anhydrous ethanol (1:1, v/v) until the leaf turned white. The absorbance of the extract was measured at 645, 663, and 440 nm using a SHIMADZU UV-1780 spectrophotometer, and the contents of Chl a, Chl b, and Car were calculated accordingly.

### 2.7. RNA sequencing and data analysis

Total RNA was isolated from Arabidopsis seedlings 7 days after treatment using the TRIzol reagent (Invitrogen, Carlsbad, CA, USA) according to the manufacturer’s protocol. RNA concentration, purity, and integrity were assessed using a NanoDrop 2000 spectrophotometer (Thermo Fisher Scientific, Waltham, MA, USA) and an Agilent 2100 Bioanalyzer (Agilent Technologies, Santa Clara, CA, USA). RNA-seq libraries were prepared from 1 μg of total RNA using the Illumina TruSeq RNA Library Prep Kit (Illumina, San Diego, CA, USA) and sequenced on an Illumina NovaSeq 6000 platform (2 × 150 bp) at Personalbio. (Shanghai, China).

After removing adapters and low-quality reads, clean reads were mapped to the Arabidopsis reference genome (TAIR10) using HISAT2 [21]. Transcripts were assembled using StringTie (v2.1.4, https://ccb.jhu.edu/software/stringtie/) [22], and gene expression levels were quantified using featureCounts (v2.0.1, http://subread.sourceforge.net/) [23] and normalized as reads per million (RPM) and transcripts per million (TPM). Differentially expressed genes (DEGs) were identified using a threshold of |log2(fold change)| > 1 and Benjamini-Hochberg adjusted *p* < 0.05. Weighted gene co-expression network analysis (WGCNA) was performed on the DEGs using the WGCNA package (v1.68) in R [24]. A soft thresholding power of 16 was chosen to construct a scale-free network. Modules were identified with a minimum module size of 30 and a cut height of 0.25. Hub genes were defined as the top 150 genes with the highest Pearson correlation coefficients. The co-expression network was visualized using Cytoscape (v3.8.0) [25].

Gene Ontology (GO) and Kyoto Encyclopedia of Genes and Genomes (KEGG) pathway enrichment analyses of DEGs were performed using the KOBAS 3.0 platform [26]. Heatmaps were generated using TBtools [27].

### 2.8. Statistical Analysis

All experiments were performed with at least three biological replicates. Data are presented as means ± standard errors (SE). One-way analysis of variance (ANOVA) followed by the least significant difference (LSD) test was performed using SPSS (v22.0) to determine significant differences among treatments at p < 0.05. Graphs were created using Graphpad prism v9.

## 3. Results

### 3.1 Differential effects of PCZ and TFS on choy sum growth and development

To investigate the effects of PCZ and TFS on the growth and development of *B. rapa*, we treated the flowering choy sum plants with 0.15 mM PCZ, 0.01 μM TFS, or a combination of both (PCZ+TFS) at the heading stage. Morphological changes were recorded at 0, 3, 5, and 7 days after treatment (DAT).

Compared with the mock-treated control, PCZ application significantly inhibited the vegetative growth of choy sum, resulting in a compact and stunted architecture. The average plant height was reduced by 41.5% at 7 DAT under PCZ treatment (Figure 1A B and 1D, Table S1). Furthermore, PCZ strongly suppressed the development of the growing point, as evidenced by the reduced bolting height and delayed flowering time. The bolting height of PCZ-treated plants was only 58.5% of that in the control group at 7 DAT (Figure 1D). The flower buds in PCZ-treated plants did not appear until 7 DAT, whereas the control plants started flowering within 3 DAT (Figure 1A and B). Besides, the newly developed leaves near the flower buds of PCZ-treated plants exhibited a crinkled and dark green phenotype (Figure 1C).

**Fig. 1.**
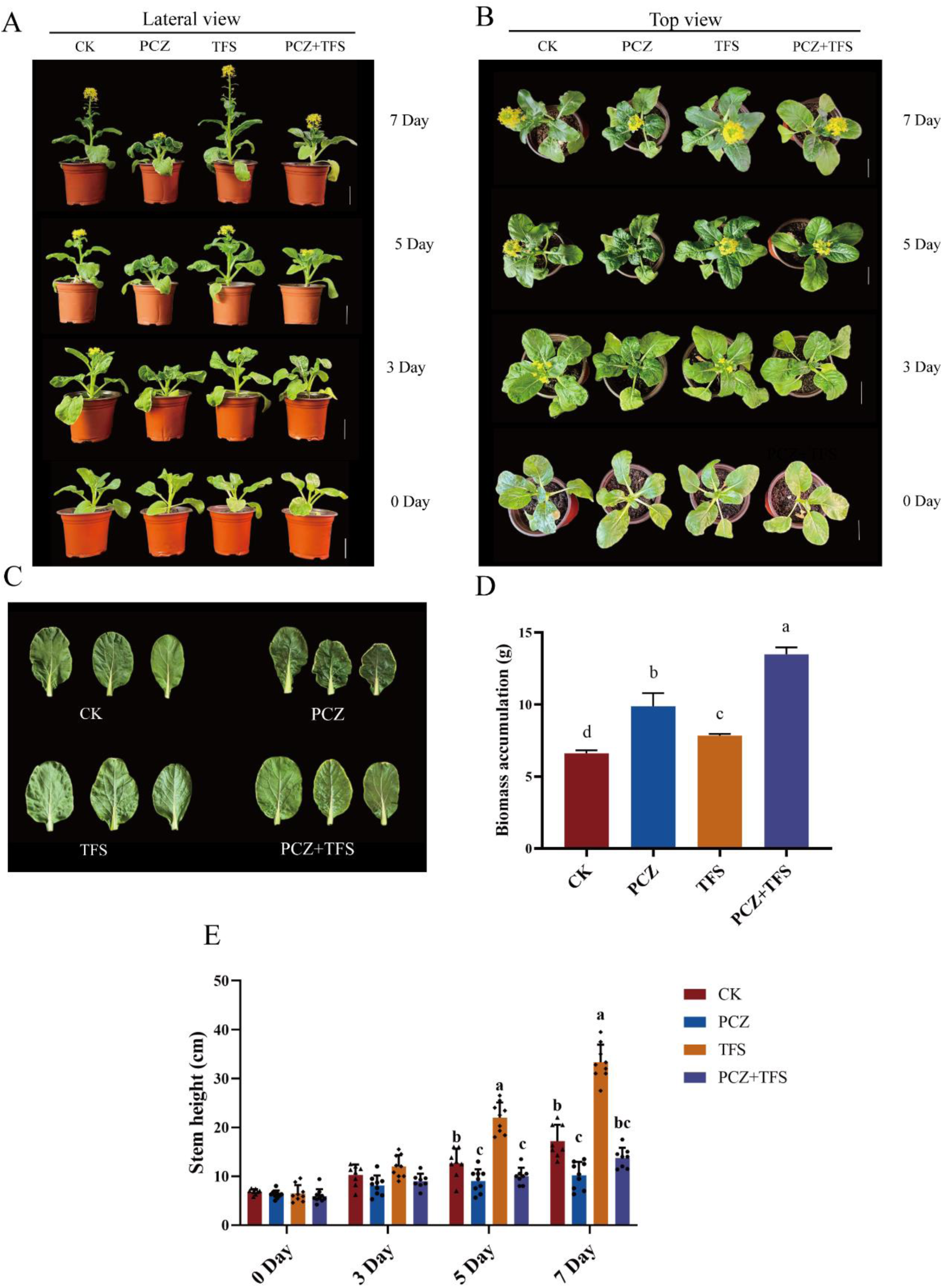
Phenotypic of *Brassica rapa* following treatment with PCZ (Propiconazole), TFS (Tianfengsu) and their combinations. (A) Growth of plants at 0, 3, 5, and 7 days after the chemical treatment. Scale bar =5 cm. (B) Leaf morphology after 7 days of treatment period. (C) Measurements of stem height over a period of 7 days following the treatments. (D) Comparison of plant weight among the different treatments. The concentrations of PCZ and TFS utilized for spraying were 0.01 μM and 0.15 mM, respectively. For the combined treatment of PCZ and TFS, a mixture was prepared by combining 0.01 μM PCZ with 0.15 mM TFS at a ratio of 1:3 (v/v). A total volume of 10 ml from each solution was applied to the plant using a miniature sprayer until both the leaves and main stem were uniformly covered with fine droplets. Stars above each bar indicates significant differences compared to the mock treatment. Data are mean ± standard deviation, n = 10 (\**P* < 0.05, \*\**P* < 0.01, \*\*\**P* < 0.001; one-way ANOVA).

In contrast to PCZ, TFS treatment promoted bolting and flowering in choy sum. The flower buds were clearly visible in TFS-treated plants at 3 DAT (Figure 1A and B). Quantitative analysis showed that the bolting height of TFS-treated plants was 1.94-fold higher than that of the control at 7 DAT (Figure 1D). To further evaluate the effects of PCZ and TFS on leaf development, we measured the leaf area of the largest rosette leaves (deformed leaves, DL) and the newly developed leaves subtending the flower buds (Qi Kou leaves, QKL). TFS treatment significantly increased the leaf area of both DL and QKL, resulting in a loose and enlarged plant architecture (Figure 1E, Table S1). Interestingly, co-application of TFS largely rescued the inhibitory effects of PCZ on choy sum growth. When treated with both PCZ and TFS, the plants showed increased height and bolting height compared to PCZ treatment alone (Figure 1D and 1E, Table S1). Moreover, TFS restored the flat and dark green appearance of the newly developed leaves in PCZ-treated plants (Figure 1C). The flowering time of PCZ+TFS-treated plants was comparable to that of the control (Figure 1A and 1B). It is worth noting that both PCZ and TFS treatments promoted biomass accumulation in choy sum, The highest biomass was observed under PCZ+TFS co-treatment, followed by PCZ and TFS single treatments (Figure 1E).

Taken together, these results suggested that PCZ and TFS treatments have opposite effects on the growth and development of flowering choy sum. PCZ inhibits vegetative growth and reproductive transition by suppressing bolting and delaying flowering, whereas TFS promotes bolting and flowering. Co-application of TFS can largely alleviate the inhibitory effects of PCZ on choy sum growth.

### 3.2 PCZ and TFS differentially enhance the flavor and nutritional quality of choy sum

Soluble sugars, soluble proteins, vitamin C, photosynthetic pigments and fatty acids are key determinants of vegetable flavor and nutritional quality [28–30]. To elucidate the effects of PCZ and TFS treatments on choy sum quality, we quantified the contents of these compounds in the leaves of treated and control plants at 7 days after treatment (DAT).

PCZ treatment significantly enhanced the accumulation of flavor-related compounds in choy sum. The contents of soluble sugars, soluble proteins, and vitamin C were increased by 25.08%, 13.68%, and 39.18%, respectively, compared to the mock control (Figure 2A-C). Similarly, TFS treatment also promoted the flavor quality of choy sum, with the contents of soluble sugars, soluble proteins, and vitamin C being increased by 74.36%, 20.71%, and 21.72%, respectively (Figure 2A-C). Notably, the combination of PCZ and TFS (PCZ+TFS) showed the strongest effects on enhancing the flavor compounds, followed by TFS and PCZ single treatments (Figure 2A-C). These results suggest that PCZ and TFS treatments differentially modulate the carbon and nitrogen metabolism in choy sum, leading to the enhanced biosynthesis and accumulation of sugars, proteins, and vitamin C [31,32].

**Fig 2:**
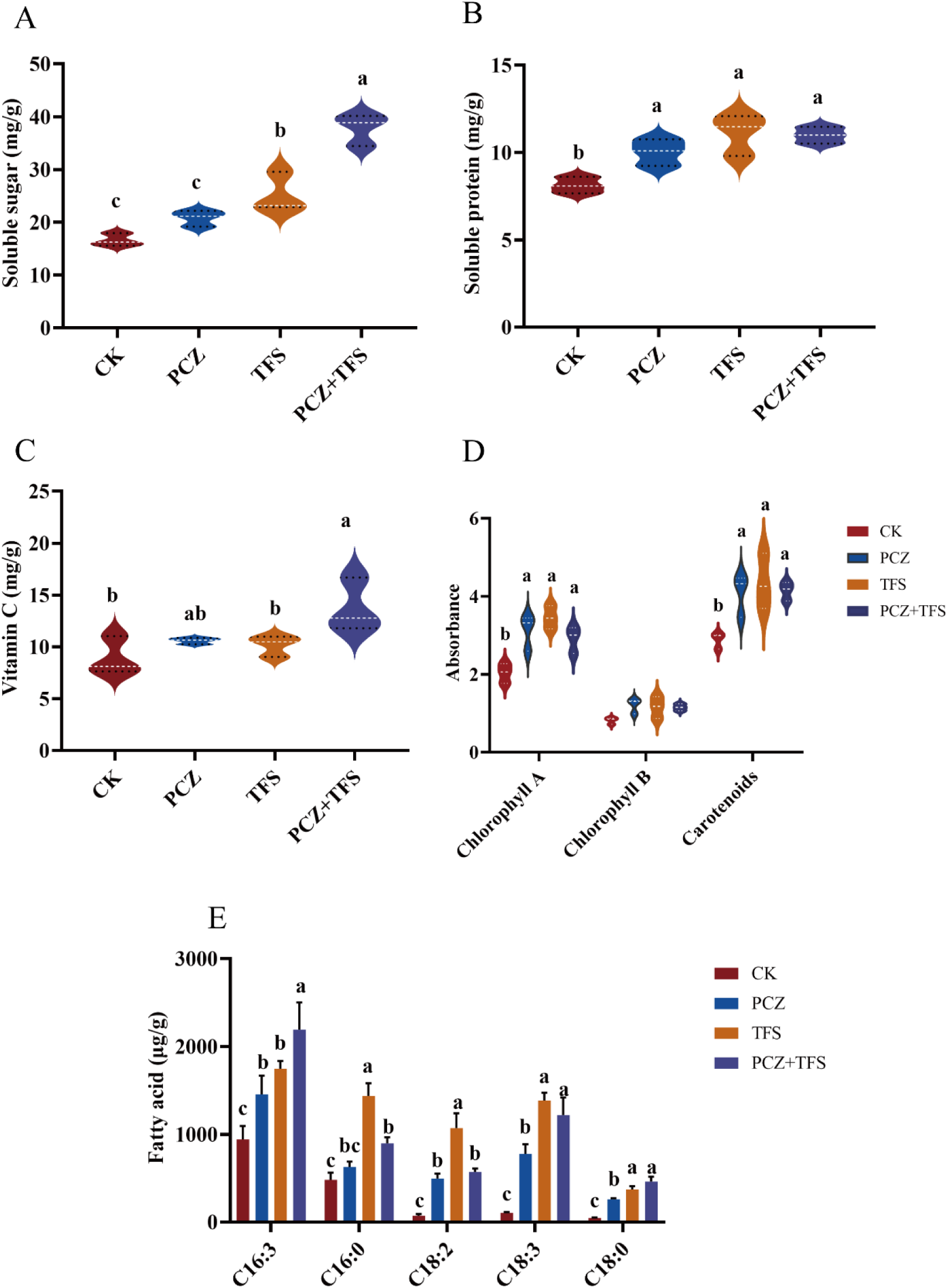
Changes of quality-related indicators in *B. rapa* in responding to treatments of PCZ, TFS, and the combination of them. (A) Soluble Sugar content, (B) Soluble Protein content, (C) Vitamin C content, (D) Photosynthetic Pigment content, and (E) Fatty Acid content (C16:3 hexadecatrienoic acid, C16:0 Palmitic acid, C18:2 Linoleic acid, C18:3 Alpha-linolenic acid, C18:0 Stearic acid). Stars indicates significant differences compared to the mock treatment. Data are mean ± standard deviation, n = 10 (\**P* < 0.05, \*\**P* < 0.01, \*\*\**P* < 0.001; one-way ANOVA).

In addition to flavor, the nutritional quality of choy sum was also greatly affected by PCZ and TFS treatments. Photosynthetic pigments, including chlorophylls and carotenoids, are important nutrients with antioxidant activities [33,34]. Both PCZ and TFS significantly increased the contents of chlorophyll a and carotenoids, while the change in chlorophyll b was not statistically significant (Figure 2D). However, the co-application of PCZ and TFS did not show a synergistic effect on the accumulation of chlorophyll a and carotenoids compared to individual treatments (Figure 2D). These results suggest that while PCZ and TFS can enhance the photosynthetic efficiency and antioxidant capacity of choy sum, their combined application may not provide additional benefits over individual treatments under the tested conditions.

Unsaturated fatty acids, such as linoleic acid (C18:2) and α-linolenic acid (C18:3), are essential nutrients that are beneficial for human health [35,36]. Fatty acid profiling revealed that PCZ and TFS treatments significantly altered the composition and contents of fatty acids in choy sum leaves (Figure 2E). The contents of several fatty acids, including hexadecatrienoic acid (C16:3), palmitic acid (C16:0), C18:2, C18:3, and stearic acid (C18:0), were increased to varying degrees under PCZ and TFS treatments. Strikingly, the accumulation of C18:3 was most strongly induced by TFS (13.2-fold), followed by PCZ+TFS (11.6-fold) and PCZ (7.4-fold), compared to the control (Figure 2E). These results indicate that PCZ and TFS treatments, especially their combination, can effectively enhance the nutritional value of choy sum by increasing the levels of health-beneficial unsaturated fatty acids.

In conclusion, our study demonstrates that PCZ and TFS treatments differentially enhance the flavor and nutritional quality of choy sum by modulating the biosynthesis and accumulation of soluble sugars, soluble proteins, vitamin C, photosynthetic pigments, and unsaturated fatty acids. The combination of PCZ and TFS exhibits the most pronounced effects on improving the overall quality of choy sum. These findings provide valuable insights into the potential application of PCZ and TFS as plant growth regulators for enhancing the flavor and nutritional quality of leafy vegetables. To gain a better understanding of the divergent impacts of PCZ and TFS on choy sum metabolism and quality development, further investigations are required to elucidate the underlying molecular mechanisms.

### 3.3 PCZ and TFS exhibit opposite effects on *Arabidopsis* rosette growth

To investigate whether PCZ and TFS treatments could produce similar effects on the growth and development of *A. thaliana* as observed in choy sum, we conducted a phenotypic analysis on the rosette leaves of treated and control plants. *Arabidopsis* seeds were germinated on 1/2 MS medium, and the seedlings were transplanted to soil at 7 days after germination (DAG). PCZ (0.15 mM), TFS (0.01 μM), or their combination (PCZ+TFS) were applied to the plants by foliar spray at 21 DAG, and the phenotypes were observed at 7 days after treatment (DAT).

Compared with the mock control, PCZ treatment significantly inhibited the growth of *Arabidopsis* rosette leaves (Figure 3A-I). The newly developed leaves of PCZ-treated plants exhibited a compact and stunted phenotype, with no obvious elongation of the petioles (Figure 3A-I). In contrast, TFS treatment promoted the growth of rosette leaves, resulting in a loose and expanded plant architecture (Figure 3A-I). Interestingly, co-application of TFS could largely alleviate the inhibitory effects of PCZ on rosette leaf growth, as evidenced by the relatively normal leaf size and plant architecture in PCZ+TFS-treated plants (Figure 3A-I).

**Fig 3.**
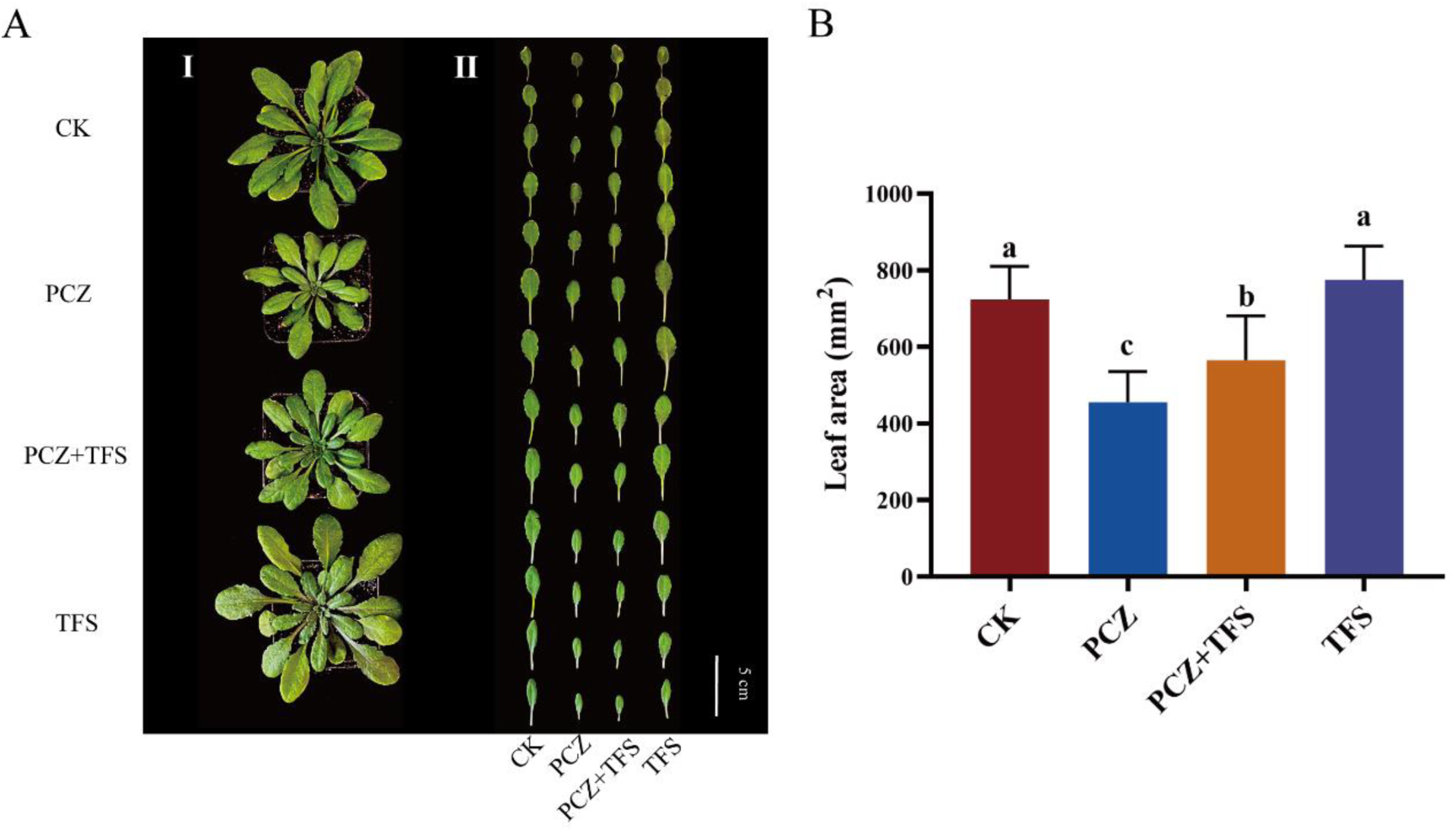
Phenotypic of *Arabidopsis thaliana* following treatment with PCZ, TFS and their combinations. (A) Growth of plants (Ⅰ) and leaf (Ⅱ) at 7 days after the chemical treatment. Scale bar =5 cm. (B) Comparison of leaf area among the different treatments. The concentrations of PCZ and TFS used for irrigation were 0.01 μM and 0.15 mM, respectively. For combined treatment of PCZ and TFS, a mixture was prepared by mixing 0.01 μM PCZ with 0.15 mM TFS in a 1:3 (v/v) ratio. A total volume of 20 ml from each solution was applied to the plant. Leaf area is expressed as length times width. Stars above each bar indicates significant differences compared to the mock treatment. Data are mean ± standard deviation, n = 10 (*\*P < 0.05, **P < 0.01, ***P < 0.001*; one-way ANOVA).

To quantify the effects of PCZ and TFS treatments on rosette leaf growth, we measured the leaf area of the treated and control plants at 7 DAT. The average leaf area of PCZ-treated plants was only 64% of that in the mock control, while the leaf area of PCZ+TFS-treated plants was 79% of the control (Figure 3A-II and 3B). These results demonstrate that PCZ inhibits the growth of *Arabidopsis* rosette leaves, and TFS can partially rescue the PCZ-induced growth defects.

Taken together, our phenotypic analysis reveals that PCZ and TFS treatments have opposite effects on the growth of *Arabidopsis* rosette leaves, similar to their differential effects on choy sum growth. PCZ suppresses rosette leaf growth and development, whereas TFS promotes leaf expansion. Co-application of TFS can alleviate the inhibitory effects of PCZ on rosette leaf growth. These findings suggest that the regulatory roles of PCZ and TFS in plant growth and development are conserved between *Arabidopsis* and *Brassica* crops, providing a valuable model system for further mechanistic studies.

### 3.4 PCZ and TFS treatments induce differential gene expression in *Arabidopsis*

To further investigate the molecular mechanisms underlying the differential effects of PCZ and TFS on plant growth and development, we performed transcriptome sequencing (RNA-seq) analysis on the rosette leaves of *Arabidopsis* plants treated with 0.15 mM PCZ, 0.01 μM TFS, or their combination (PCZ+TFS) at 7 days after treatment (DAT). The mock-treated plants served as the control (CK).

RNA-seq analysis identified a total of 2,486, 1,789, and 3,242 differentially expressed genes (DEGs) in PCZ, TFS, and PCZ+TFS treated plants, respectively, compared to the CK (Figure S1 and S2). Among them, 1,328 (53.4%), 964 (53.9%), and 1,832 (56.5%) DEGs were upregulated, while 1,158 (46.6%), 825 (46.1%), and 1,410 (43.5%) DEGs were downregulated in the PCZ, TFS, and PCZ+TFS groups, respectively (Figure S2). Venn diagram analysis revealed that 612 DEGs were commonly regulated by all three treatments. Additionally, PCZ specifically induced 1,026 DEGs, TFS specifically induced 339 DEGs, and the combination of PCZ and TFS specifically induced 1,363 DEGs (Figure S3A). Principal component analysis (PCA) and hierarchical clustering analysis showed that the transcriptome profiles of PCZ and PCZ+TFS treated plants were more similar to each other than to those of TFS and CK groups (Figure S3B and S3C), indicating that PCZ had a dominant effect on the global gene expression pattern in *Arabidopsis* rosette leaves.

Gene Ontology (GO) enrichment analysis revealed that the DEGs induced by PCZ and PCZ+TFS were significantly enriched in the biological processes related to hormone signaling (brassinosteroid and auxin), cell wall organization, photosynthesis, and secondary metabolite biosynthesis (phenylpropanoid and flavonoid) (Figure S4A and S4C). In contrast, the TFS-induced DEGs were mainly involved in the biological processes of cell growth, cell cycle, DNA replication, and meristem development (Figure S4B). These results suggest that PCZ and TFS treatments differentially regulate the transcriptional programs associated with hormone signaling, cell wall remodeling, and secondary metabolism in *Arabidopsis*.

Kyoto Encyclopedia of Genes and Genomes (KEGG) pathway enrichment analysis revealed that PCZ and PCZ+TFS treatments significantly affected several metabolic pathways closely related to the biosynthesis and accumulation of flavor compounds in plants, including starch and sucrose metabolism, glycolysis, citrate cycle, amino acid metabolism, and biosynthesis of secondary metabolites (Figure 4A and 4C). Notably, the BR biosynthesis pathway was also enriched among the PCZ-responsive genes, with several BR biosynthetic genes being upregulated while the BR catabolic gene BAS1 was induced (Figure 5), suggesting a feedback regulation of BR homeostasis in response to PCZ-mediated BR deficiency [37]. The changes in BR level and signaling may have a profound impact on the downstream metabolic pathways and the accumulation of flavor-related compounds. For instance, the PCZ-induced enrichment of starch and sucrose metabolism, citrate cycle, and amino acid metabolic pathways (Figure 4A) coincided with the increased contents of soluble sugars, soluble proteins, and organic acids in PCZ-treated choy sum leaves (Figure 2A-C), implying that BR deficiency may reprogram carbon and nitrogen metabolism to favor the accumulation of these flavor compounds.

**Fig 4.**
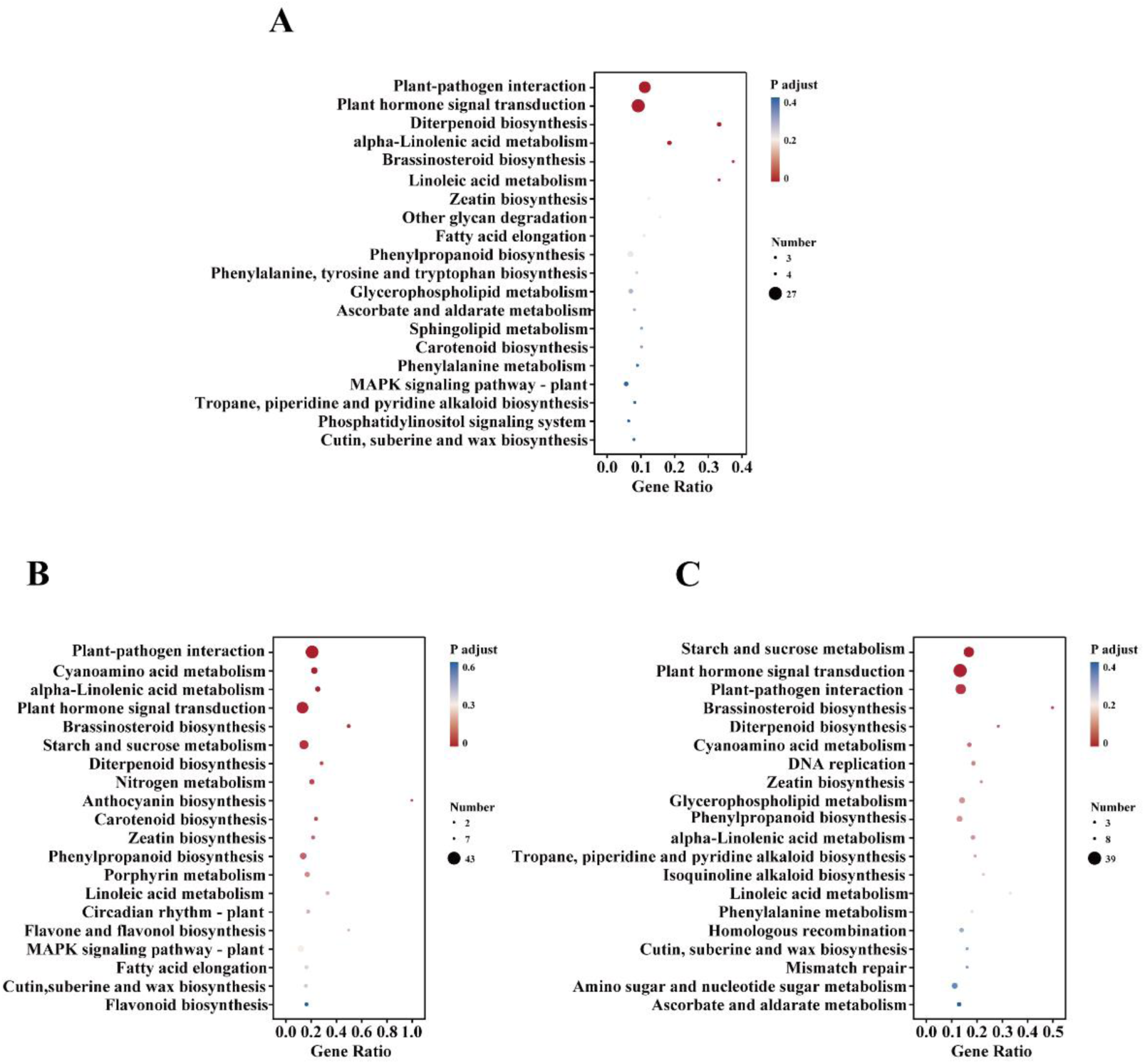
KEGG pathway analysis in leaves of *Arabidopsis thaliana* exposed to PCZ, TFS, and the combination of them. (A) PCZ, (B) TFS and (C) PCZ+TFS. Samples were collected over a period of 7 days following the treatments with 0.01 μM PCZ, or 0.15 mM TFS, or a mixture was prepared by combining 0.01 μM PCZ with 0.15 mM TFS at a ratio of 1:3 (v/v). *p*-value represents the statistical test value (*t*-test) of the difference in gene expression level. The gene-rich factor was calculated as the gene ratio of the differential expression genes (DEGs) assigned to the KEGG pathway/total DEGs.

**Fig. 5.**
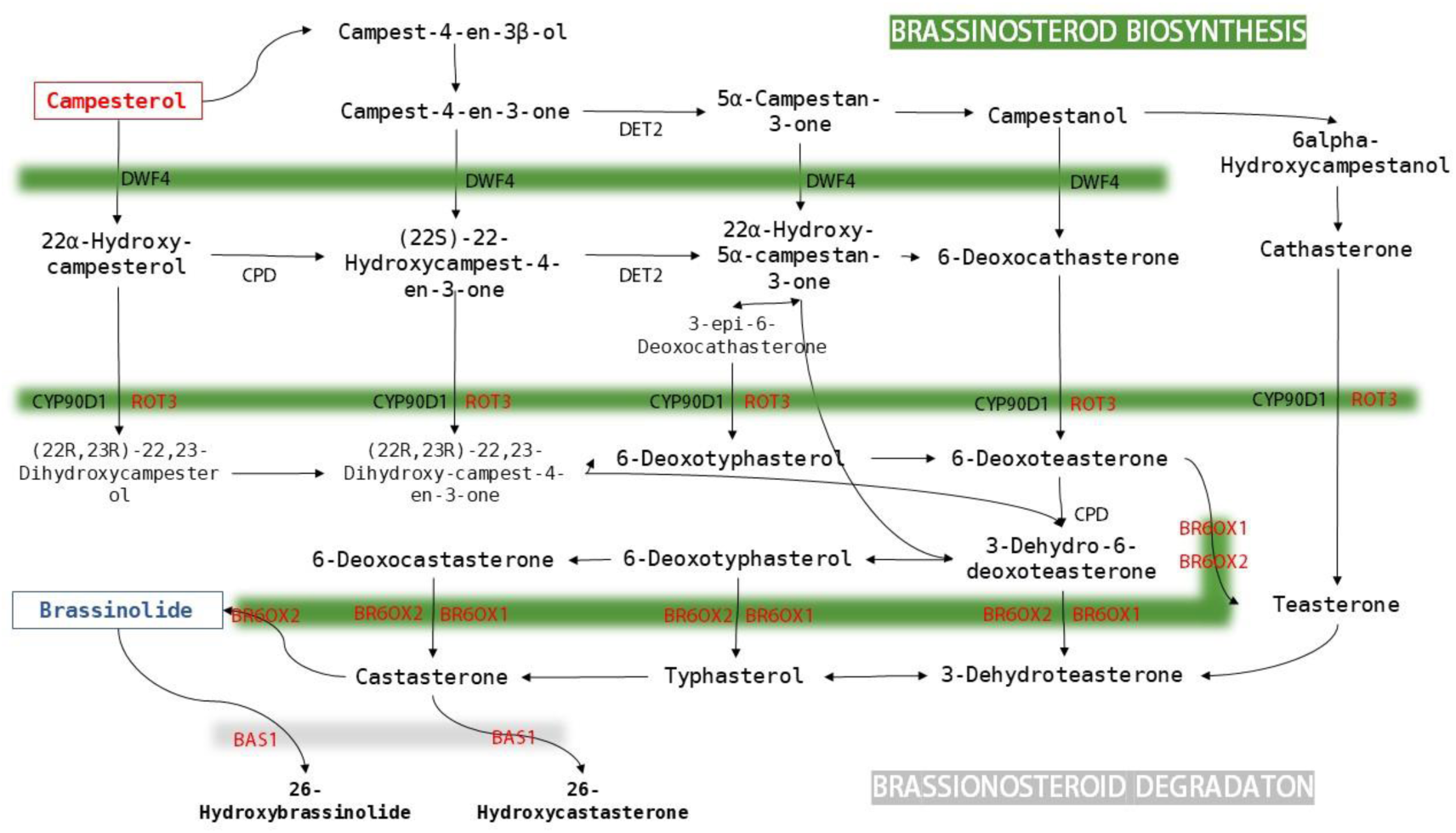
Brassinosteroid biosynthesis pathway in *Arabidopsis thaliana* PCZ treatment. Red indicates differential up-regulation of related genes/metabolites, and blue indicates differential down-regulation of related genes/metabolites.

Interestingly, TFS co-treatment further enhanced the enrichment of starch and sucrose metabolism, amino acid metabolism, and biosynthesis of secondary metabolites (Figure 4C), which may account for the more prominent increase in soluble sugars, soluble proteins, and pigments in the PCZ+TFS group compared to PCZ treatment alone (Figure 2A-D). Moreover, TFS treatment specifically enriched the pathways related to the biosynthesis of carotenoids and chlorophylls (Figure 4B), in line with the significant increase in chlorophyll a and carotenoid contents under TFS treatment (Figure 2D). These findings suggest that TFS may partially counteract the effects of PCZ on BR signaling and primary metabolism while synergistically promoting the accumulation of secondary metabolites and pigments [38].

Our transcriptome analysis reveals that PCZ and TFS treatments differentially modulate the expression of genes involved in hormone signaling, primary metabolism, and secondary metabolism in *Arabidopsis* rosette leaves. The PCZ-induced BR deficiency appears to enhance the accumulation of soluble sugars, amino acids, and organic acids by reprogramming the carbon and nitrogen metabolic pathways, whereas TFS may rescue the PCZ effects on primary metabolism and further promote the biosynthesis of pigments and secondary metabolites. The synergistic actions of PCZ and TFS on multiple metabolic pathways may contribute to the improved flavor and nutritional quality of choy sum. To further dissect the gene regulatory networks underlying these metabolic changes, we performed weighted gene co-expression network analysis (WGCNA) on the DEGs.

### 3.5 Identification of Key DEGs involved in drug response by WGCNA

To elucidate the coordinated regulation of genes in response to drug treatments, we performed weighted gene co-expression network analysis (WGCNA) on the differentially expressed genes (DEGs) from different treatment groups. DEGs with FPKM > 1 were clustered into seven distinct modules (Figure 6A), with the turquoise module containing the largest number of DEGs (477) and the grey module having the fewest (5). The module-trait relationship heatmap revealed that genes in the turquoise module positively correlated with the control group and negatively correlated with the drug treatment groups, while genes in the yellow module exhibited the opposite trend (Figure 6B), suggesting their involvement in the physiological and flavor-related changes in choy sum under drug treatments.

**Fig 6.**
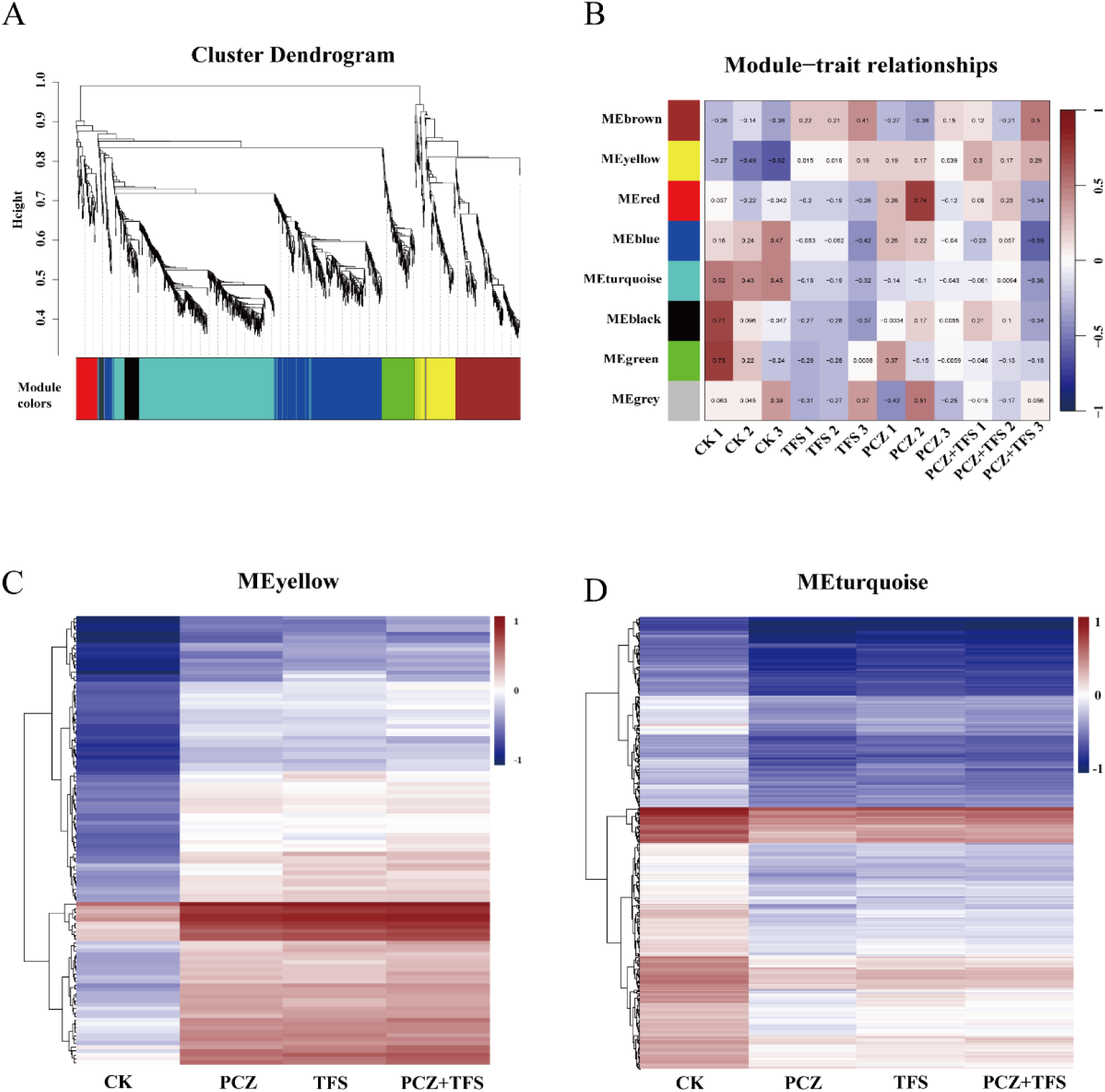
Weighted gene co-expression network analysis (WGCNA) analysis of all DEGs with FPKM >1 in leaves of *Arabidopsis thaliana* exposed to PCZ, TFS, and the combination of them. (A) Hierarchical clustering dendrogram of all DEGs. (B) The relationship among co-expression modules and different treatment groups. heatmap of genes expression of the yellow module (C) and turquoise module (D). Module-trait relationships are represented by color changes Correlation Coefficients. The FPKM of DEGs is represented by the color change from red to blue after normalization.

Hierarchical clustering analysis showed that over 90% of the genes in the turquoise module were downregulated, while more than 90% of the genes in the yellow module were upregulated following PCZ and TFS treatments (Figure 6C, D). KEGG enrichment analysis revealed that the turquoise module genes were enriched in α-linolenic acid metabolism, glycerophospholipid metabolism, starch and sucrose metabolism, and cyanogenic amino acid metabolism (Figure 7A), pathways closely related to the accumulation of flavor compounds such as soluble sugars and fatty acids. Conversely, the yellow module genes were enriched in carotenoid biosynthesis, plant hormone signal transduction, and cysteine and methionine metabolism (Figure 7B), associated with photosynthetic pigment biosynthesis and plant growth regulation.

**Fig 7.**
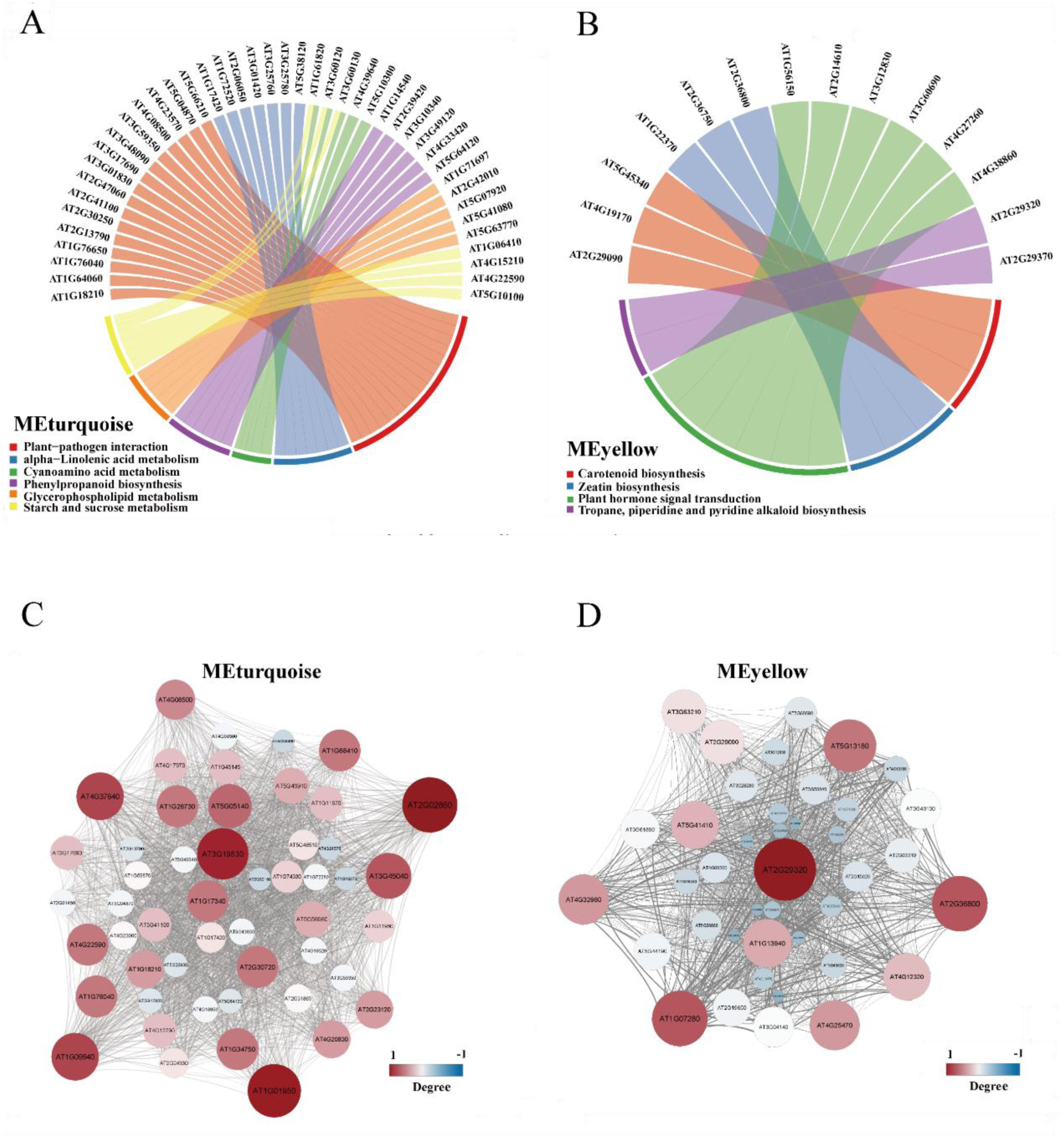
KEGG pathway enrichment of key module genes. (A) MEturquoise, (B) MEyellow. 40 Hub gene network diagrams of specific modules. (C) MEturquoise, (D) MEyellow. In network diagrams, The color change from red to blue and the size of the circle represent the degree of the gene, and the change in gene transparency in the thickness of the connecting line represents the correlation between genes.

Gene interaction network analysis identified hub genes within each module (Figure 7C and 7D). In the turquoise module, hub genes such as *SUT2*, *ARK2*, *NTMC2T5.2*, *ACA2*, and *HEMA2* were primarily involved in regulating soluble sugar accumulation, plant morphogenesis, and chlorophyll synthesis (Table S2). Their downregulation under drug treatments (Table S3) is consistent with the observed changes in flavor compounds and plant growth. In the yellow module, hub genes like *DOGT1* and *ATH1* were mainly associated with reducing BR levels and delaying flowering time (Table S2), and their upregulation (Table S3) may contribute to the dwarf and compact phenotype of choy sum under PCZ treatment.

The *SUT2* gene, encoding a sucrose transporter, plays a crucial role in controlling sucrose transport from mesophyll cells to sieve tubes [39,40]. Its downregulation under drug treatments may lead to the accumulation of soluble sugars in choy sum leaves. The *ARK2* gene, involved in tubulin formation and plant morphogenesis [41,42], and the *HEMA2* gene, encoding a key enzyme in the chlorophyll synthesis pathway [43], were also downregulated, potentially contributing to the altered plant architecture and photosynthetic pigment content. On the other hand, the upregulation of the *DOGT1* gene, which glucosylates brassinosteroids and leads to a BR-deficient phenotype when overexpressed [44], and the *ATH1* gene, a transcription factor involved in photomorphogenesis and flowering time regulation [45, 46], may be responsible for the dwarf and compact phenotype and delayed flowering observed in PCZ-treated choy sum.

## 4. Discussion

In this study, we investigated the differential effects of propiconazole (PCZ) and Tianfengsu (TFS) on the growth, development, flavor, and nutritional quality of choy sum (*B. rapa*) and *A. thaliana*. Our results demonstrate that PCZ and TFS treatments have opposite effects on plant growth and development, with PCZ inhibiting while TFS promoting these processes. Interestingly, co-application of TFS can largely rescue the inhibitory effects of PCZ on growth while synergistically enhancing the flavor and nutritional quality. Transcriptomic analysis in *A. thaliana* that PCZ and TFS differentially regulate the expression of genes involved in brassinosteroid (BR) signaling and downstream metabolic pathways, providing molecular insights into their phenotypic effects.

PCZ is known to inhibit BR biosynthesis by targeting cytochrome P450 monooxygenases, leading to BR deficiency [47,48]. Consistent with this, we observed that PCZ treatment induced BR deficiency phenotypes in both choy sum and Arabidopsis, such as dwarfism, dark green leaves, and delayed flowering [49]. Transcriptomic analysis in *A. thaliana* revealed a feedback upregulation of BR biosynthetic genes and downregulation of BR-responsive genes under PCZ treatment, further confirming the disruption of BR homeostasis [50–52]. Despite the growth inhibition, PCZ enhanced the accumulation of flavor compounds, including soluble sugars, proteins, and vitamin C in choy sum. This suggests a reprogramming of primary and secondary metabolism in response to BR deficiency, as supported by the enrichment of related pathways in PCZ-treated Arabidopsis [53,54].

In contrast, TFS treatment promoted growth and development in choy sum and *A. thaliana*, likely by activating BR signaling [58,59]. Moreover, TFS increased the contents of photosynthetic pigments and unsaturated fatty acids in choy sum, enhancing its nutritional value. The transcriptomic data revealed an upregulation of genes involved in pigment and fatty acid biosynthesis under TFS treatment [60,61], providing a molecular basis for these metabolic changes.

Remarkably, co-application of TFS could largely rescue the PCZ-induced growth inhibition while further enhancing the flavor and nutritional quality of choy sum. This suggests a fine-tuning of BR signaling and downstream metabolic pathways by the combined treatment [62,63]. The transcriptomic analysis in *A. thaliana* supports this notion by showing a restoration of BR homeostasis and a synergistic enhancement of multiple metabolic pathways under PCZ+TFS treatment.

To further dissect the gene regulatory networks underlying these differential effects, we performed weighted gene co-expression network analysis (WGCNA) and identified key modules and hub genes associated with the phenotypic and metabolic changes. For example, the SUT2 and HEMA2 genes in the turquoise module were downregulated under PCZ and TFS treatments, potentially contributing to the accumulation of soluble sugars and altered leaf morphology [32–34]. In the yellow module, the upregulation of BR catabolic gene DOGT1 and flowering repressor ATH1 may account for the dwarf phenotype and delayed flowering under PCZ treatment [37–39].

Our findings are consistent with a recent study by Song et al. [64], which examined the effects of PCZ on soybean growth and development. However, our study provides a more comprehensive analysis by investigating not only the changes in growth and development but also the alterations in flavor and nutritional components in *B. rapa*. Furthermore, we performed a deeper exploration of the molecular mechanisms by focusing on the expression changes of BR signaling pathway-related genes, confirming that the BR pathway participates in regulating the growth, development, flavor, and nutritional responses of B. rapa to PCZ and TFS treatments. These results suggest that BRs play conserved roles in regulating plant growth and development across different species, with physiological responses tightly coupled to transcriptional changes in multiple metabolic pathways [65,66]. By comparing PCZ and TFS treatments, we elucidated the mechanisms underlying their differential impacts on growth and flavor quality.

It is important to note that the combined application of PCZ and TFS achieved the best balance between growth, flavor, and nutritional quality in choy sum. However, PCZ is not registered for use on choy sum in China, and its off-label application raises concerns about food safety and agricultural sustainability. Our study provides valuable insights into the optimal concentration and application timing of PCZ for achieving the desired growth control while minimizing potential risks. Furthermore, the identification of key genes and pathways involved in BR-mediated regulation of growth and quality traits offers new targets for developing alternative strategies, such as screening for novel plant growth regulators or breeding high-quality choy sum varieties that meet both consumer demands and production requirements.

In conclusion, our study provides a comprehensive understanding of how PCZ and TFS differentially regulate plant growth, flavor, and nutritional quality through modulating BR signaling and metabolism. These findings not only advance our knowledge of the molecular mechanisms underlying vegetable quality improvement but also have important implications for guiding the safe and efficient production of high-quality choy sum. Future research should focus on determining the minimum effective and safe concentration of PCZ, establishing appropriate application intervals, and exploring alternative approaches for optimizing choy sum growth and quality, such as developing new plant growth regulators or breeding elite varieties based on the key genes and pathways identified in this study.

## 5. Conclusion

In summary, this study reveals that PCZ and TFS differentially regulate the growth, flavor, and nutritional quality of choy sum and *Arabidopsis* by modulating BR signaling and downstream metabolic pathways. PCZ inhibits growth while enhancing flavor compound accumulation, whereas TFS promotes growth and increases photosynthetic pigments and unsaturated fatty acids. Co-application of PCZ and TFS synergistically improves overall quality. Transcriptomic analysis in *A. thaliana* identifies key BR-related genes and pathways underlying these differential effects, providing novel insights into the molecular mechanisms of flavor regulation in leafy vegetables. These findings offer valuable guidance for developing strategies to improve vegetable quality through rational application of plant growth regulators and targeted breeding approaches.

## Author Contribution

Fei Lin and Hanhong Xu: Conceptualization, Resources, Supervision, Funding acquisition, Writing - review & editing. Dekang Guo: Conceptualization, Methodology, Investigation, Formal analysis, Writing - original draft. Qing Gao: Investigation, Formal analysis. Yunxue Song: Investigation. Zhicheng Liu: Plants Cultivation. Daorui Wang: conceptualization and methodology.

## Acknowledgements

This work was supported by the Jiangmen Science and Technology Commissioners Project (2023760100240008456). Generic Technique Innovation Team Construction of Modern Agriculture of Guangdong Province (2023KJ130) and Guangdong Province Key Areas R&D Plan Project (2022B0202080001). We thank the *Arabidopsis* Biological Resource Center (ABRC) for providing the *Arabidopsis thaliana* Col-0 seeds.

## Appendix A. Supplementary data

**Fig S1.**
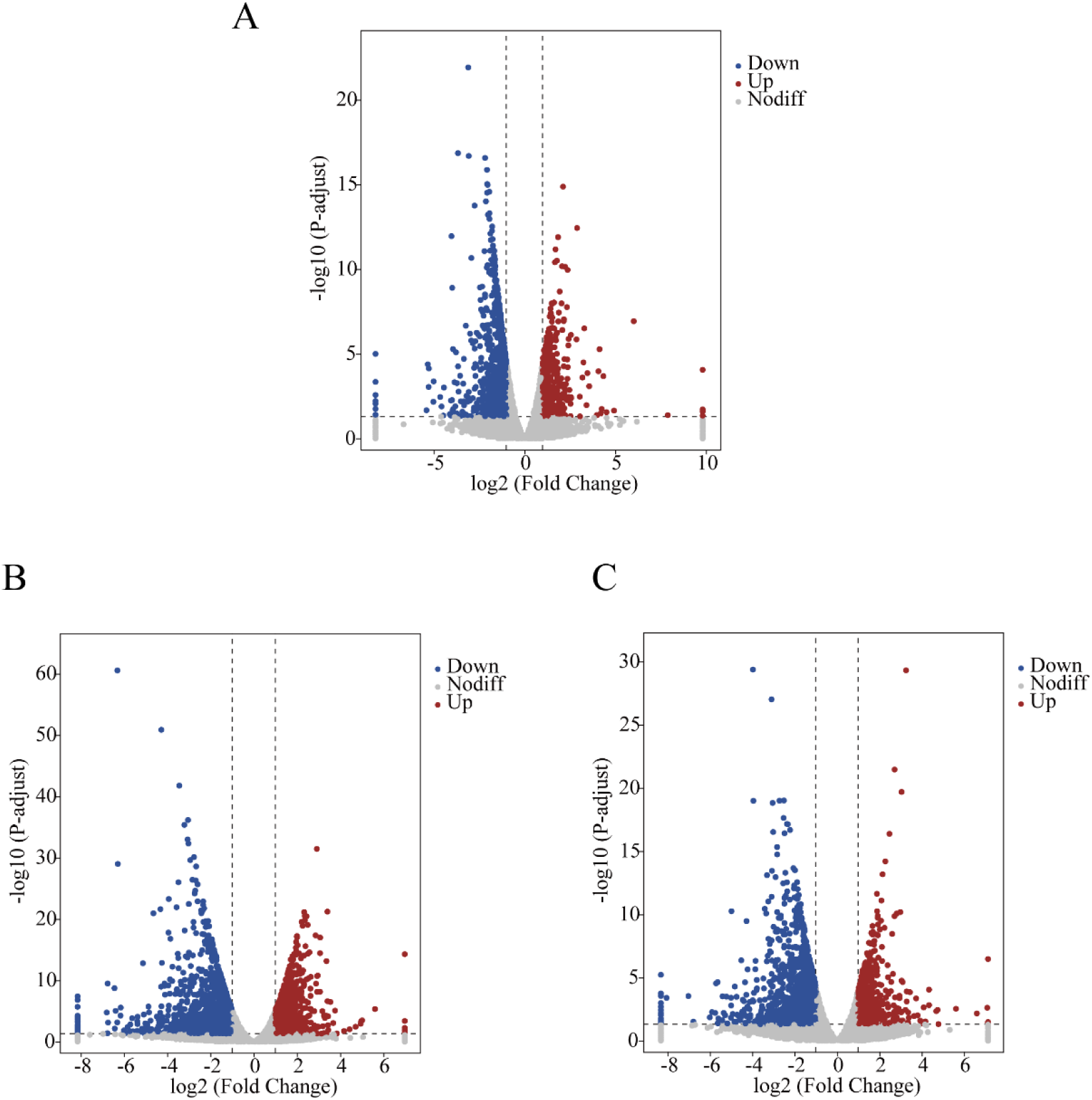
Volcano plots of DEGs in *A. thaliana* treated with PCZ (A), TFS (B), and FCZ+TFS (C) compared to the CK group (*p* < 0.05). The plots display log10 of the minimum *P*-values among the four alternative hypotheses plotted against the Log_2_FC of the fold change of DEGs, with the positive and negative values represent the up- and down-regulated of DEGs, respectively. The colors of data points indicate the differential expression categories to which a gene has been classified.

**Fig S2.**
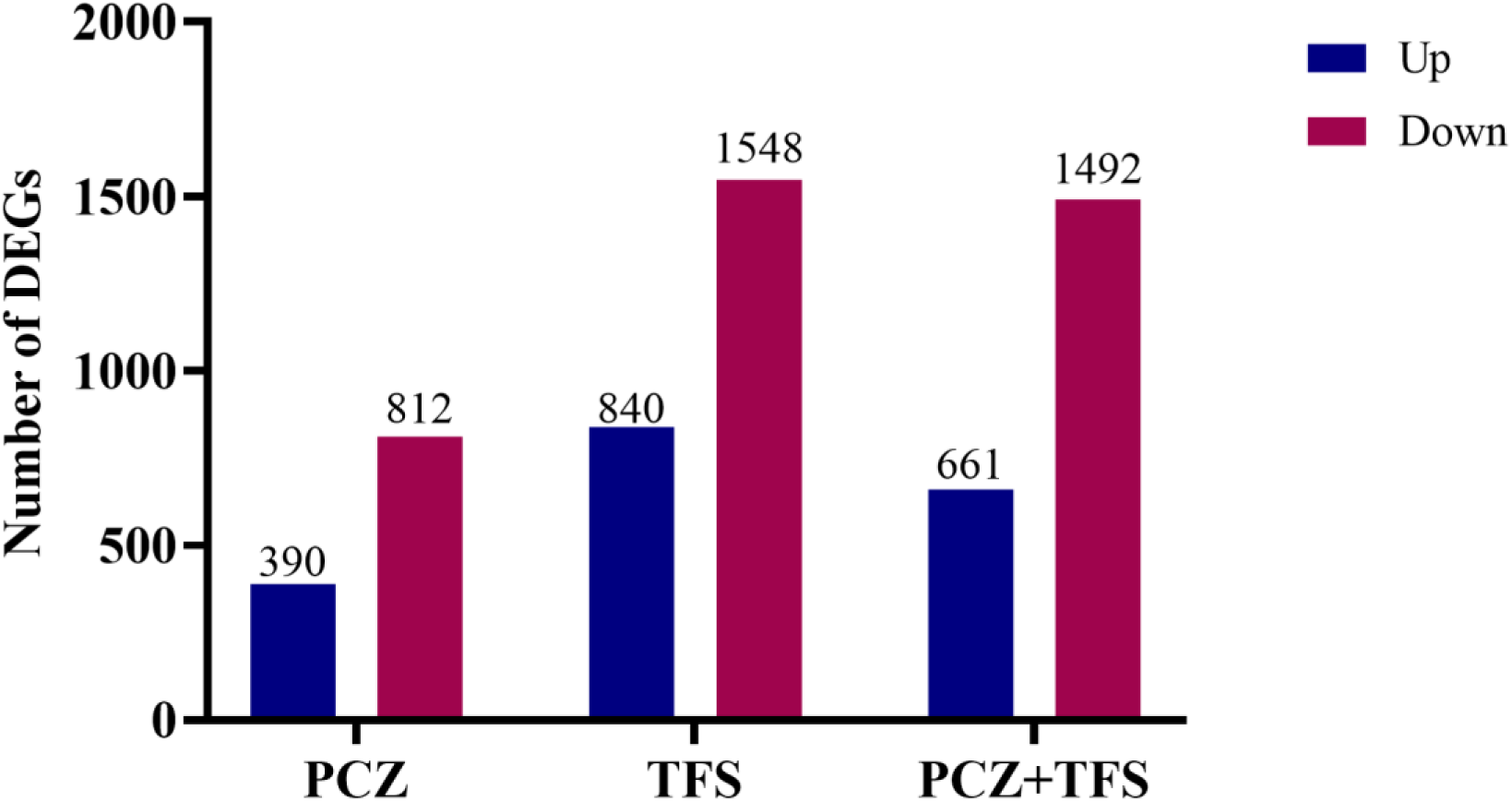
Number of differentially expressed genes (DEGs) in *A. thaliana* treated with PCZ, TFS and their combination.

**Fig S3.**
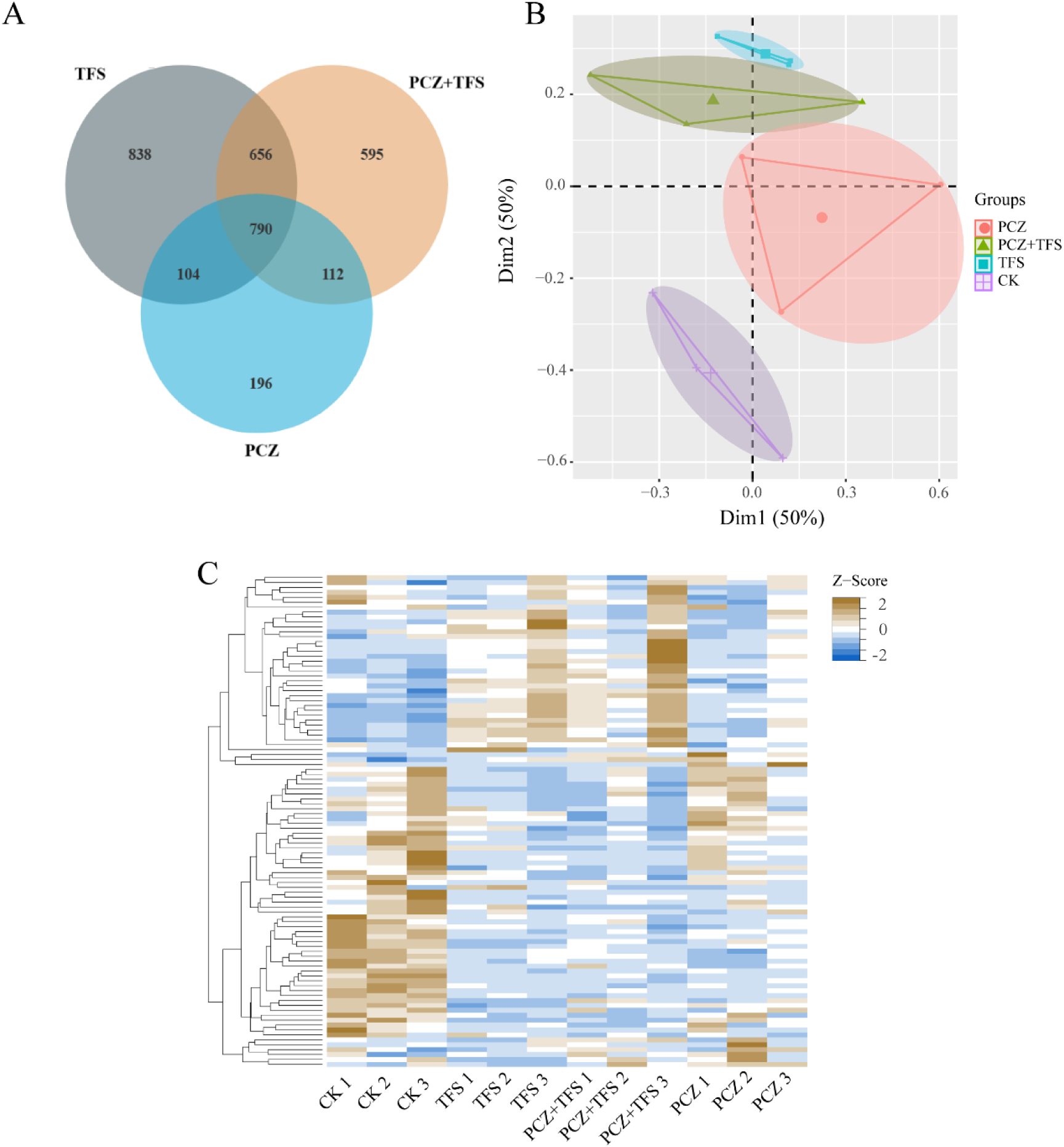
Variance and cluster analysis of *A. thaliana* under treatments of PCZ, TFS, and a mixture of them. (A) Venn diagram of DEGs. The green circle represents the comparison between CK and PCZ datasets, while the purple blue circle represents the comparison between CK and TFS datasets; the red circle represents the comparison between CK and PCZ+TFS datasets. The overlapping region of these circles indicates the shared DEGs among these datasets; (B) Principal component analysis (PCA) of DEGs. The diagram illustrates the PCZ, TFS, and PCZ+TFS groups along with the CK group. The dots of the same color represent three biological replicates within each treatment group. (C) Cluster heat map analysis of DEGs. The abscissa represents different samples, while the vertical axis represents clusters of differentially expressed genes (DEGs). The color red indicates upregulation, whereas green indicates downregulation. The legend in the heat map represents the Z-score after normalization of gene expression.

**Fig S4.**
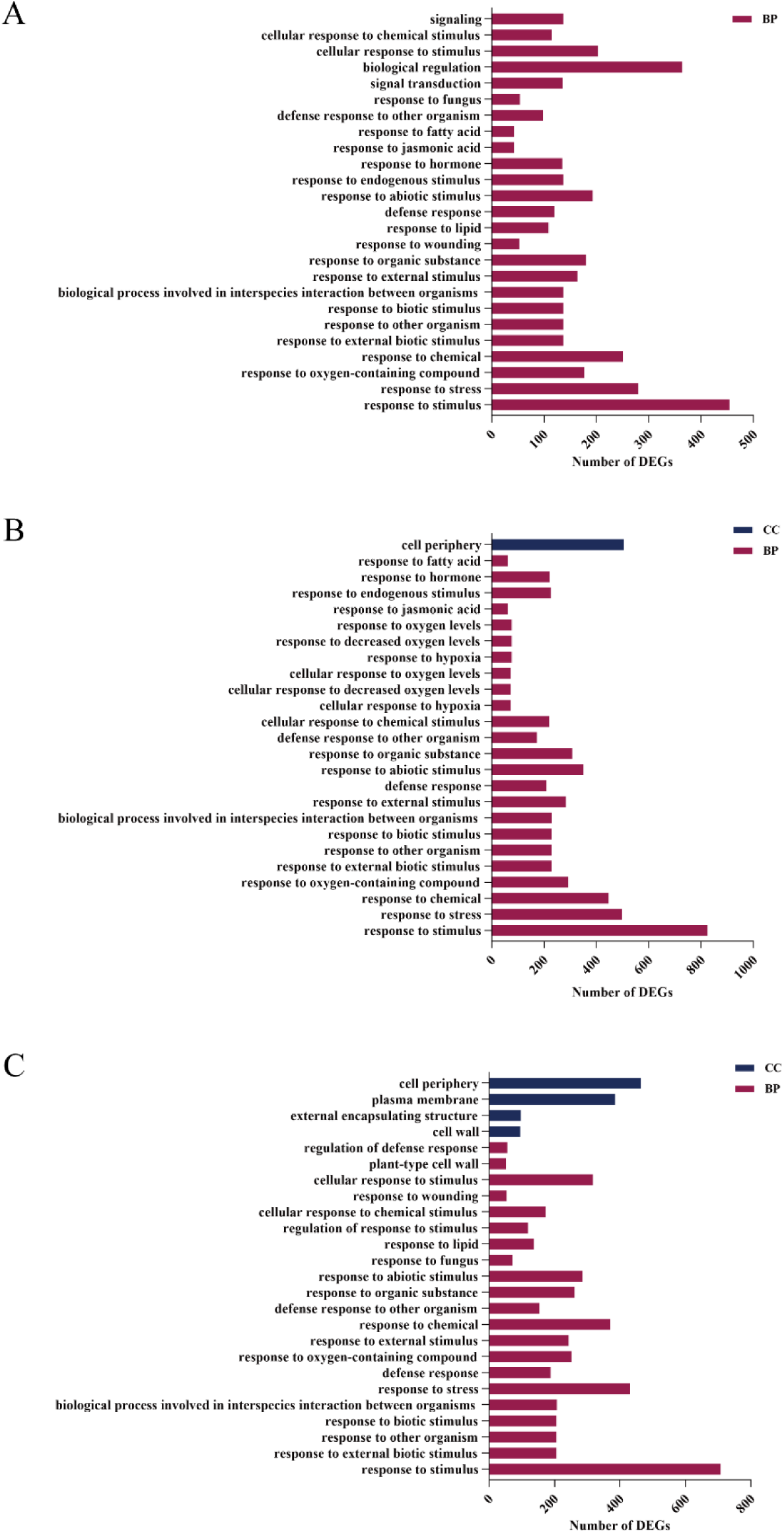
GO annotation analysis of DEGs in *A. thaliana* treated with PCZ (A), TFS (B), and the PCZ+TFS (C). The abscissa represents the GO term, while the enrichment denotes the number of differential genes associated with that term. GO terms exhibiting significant enrichment in the treatment compared to the CK group are selected for visualization on the map. The red bar column indicates DEGs enriched in biological processes (BP), and the orange bar column signifies DEGs enriched in cellular components (CC).

**Table S1.**
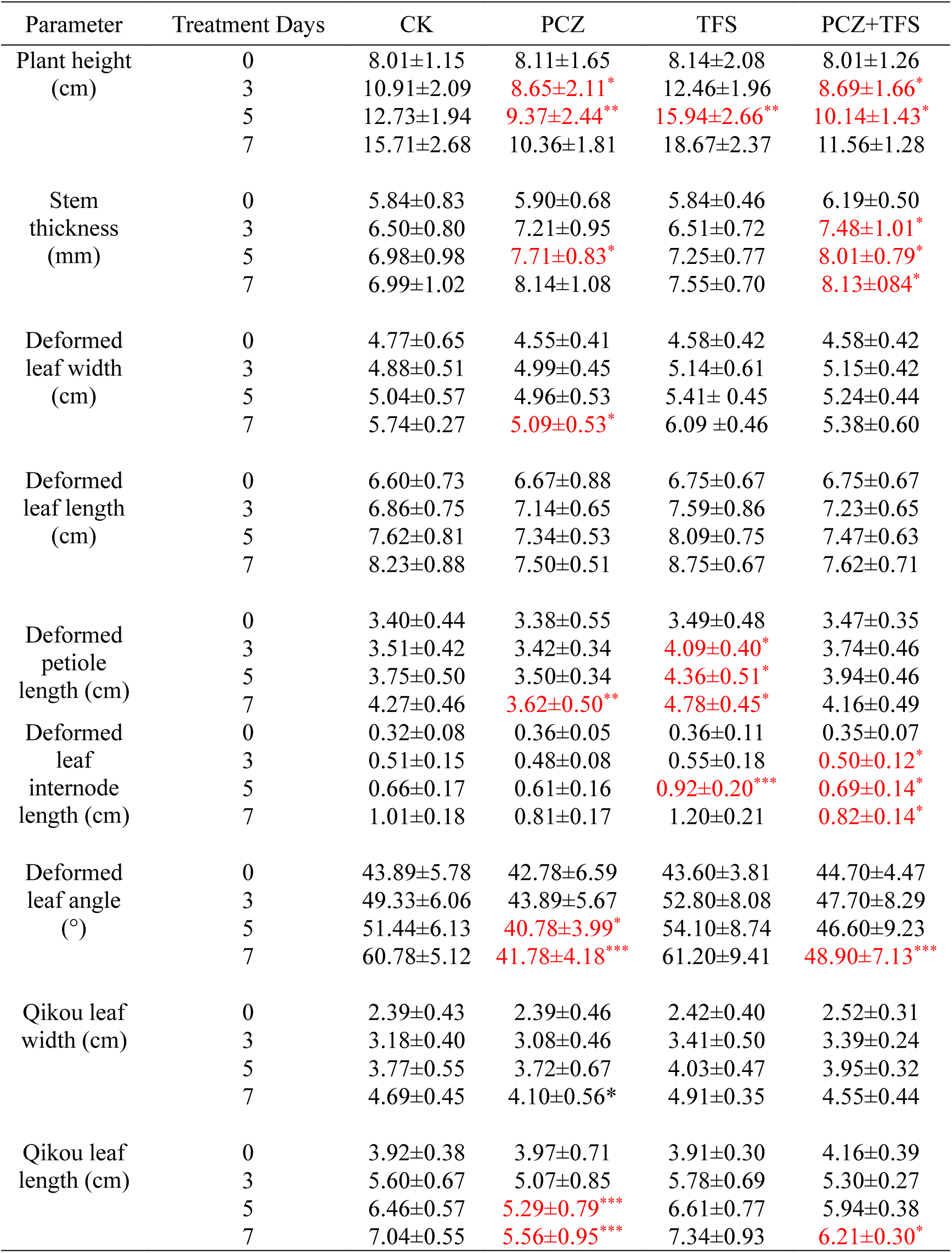
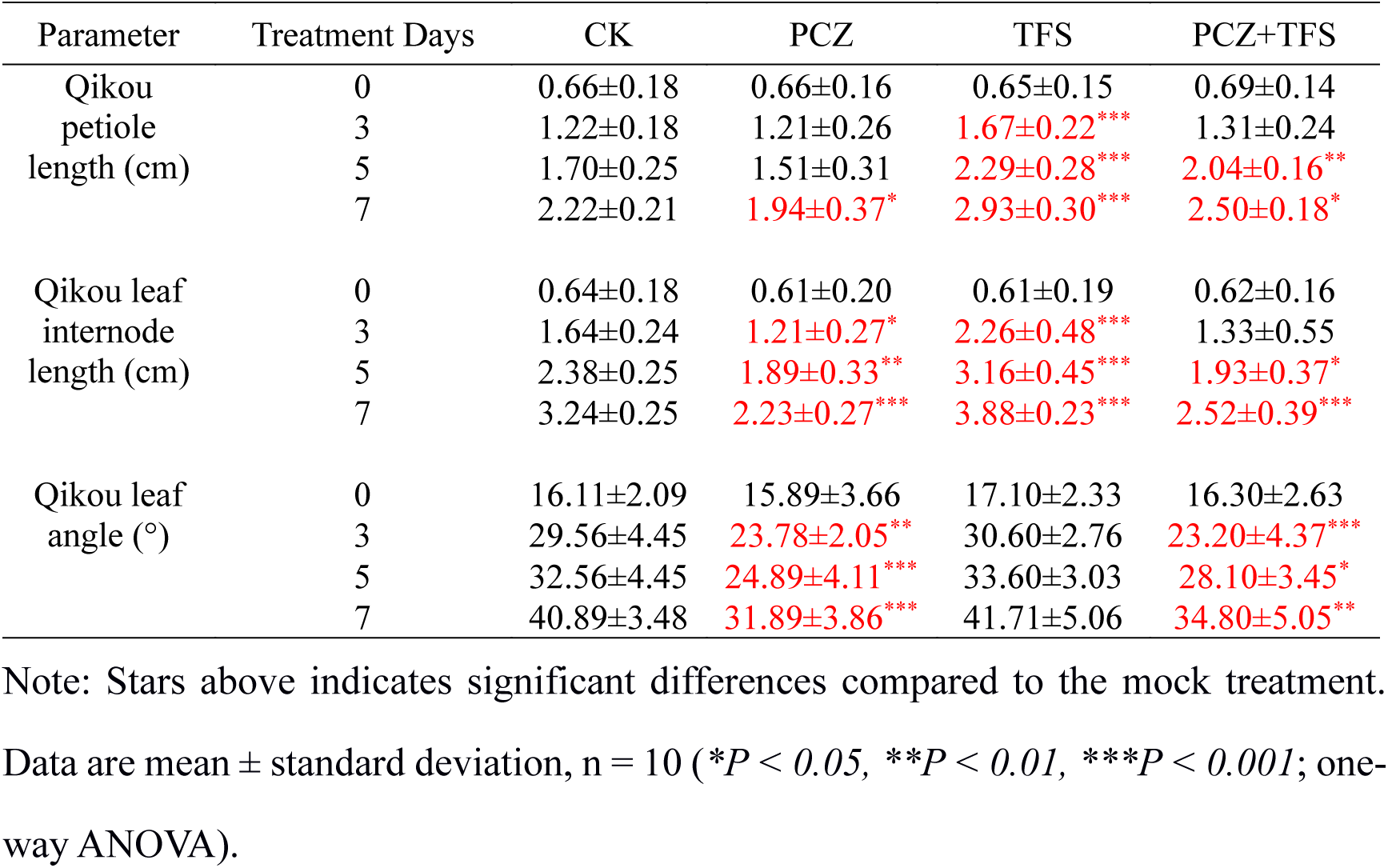
Phenotypic statistics of Brassica rapa under the combined treatment of PCZ, TFS and their combinations.

**Tabel S2.** Differential changes in differentially expressed genes in brassinosteroid-related pathways.

| Gene ID | Differential Genes | Gene name | logFC_PCZ | logFC_TFS | logFC_PT |
| --- | --- | --- | --- | --- | --- |
| AT4G36380 | Cytochrome P450 superfamily protein | ROT3 | 2.07 | 1.09 | 1.58 |
| AT5G38970 | brassinosteroid-6-oxidase 1 | BR6OX1 | 2.28 | 2.66 | 1.77 |
| AT3G30180 | brassinosteroid-6-oxidase 2 | BR6OX2 | 3.57 | 2.15 | 2.82 |
| AT3G13730 | cytochrome P450, family 90, subfamily D, polypeptide 1 | CYP90D1 | 0.97 | 0.74 | 1.07 |
| AT5G05690 | carboxypeptidase D | CPD | 0.85 | 0.2 | 0.64 |
| AT3G50660 | Cytochrome P450 superfamily protein | DWF4 | 0.22 | -0.04 | -0.12 |
| AT2G38050 | 3-oxo-5-alpha-steroid 4-dehydrogenase family protein | DET2 | -0.02 | 0.58 | 0.72 |
| AT2G26710 | Cytochrome P450 superfamily protein | BAS1 | 2.36 | 0.9 | 0.92 |

**Tabel S3.**
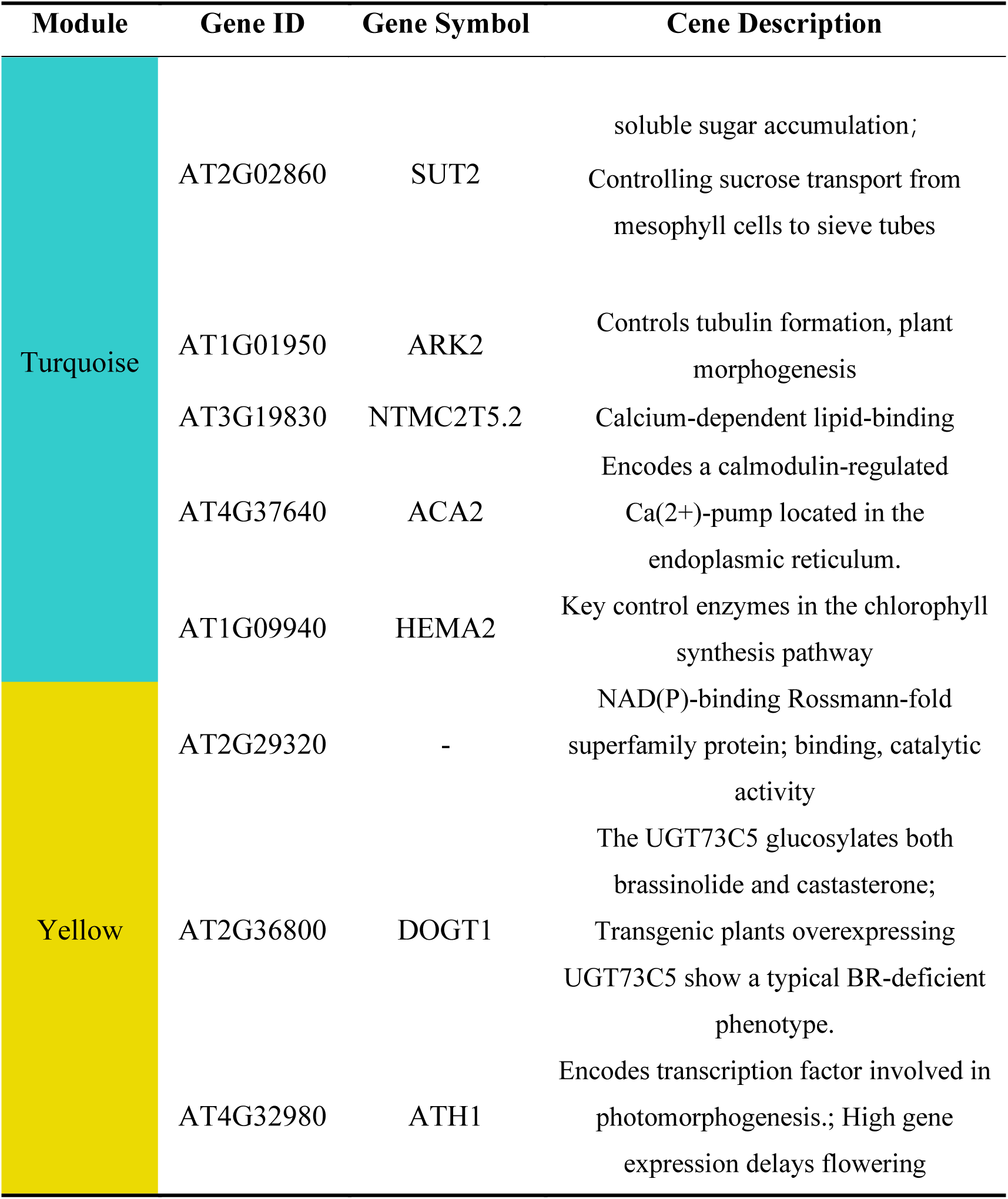
Hub gene table in turquoise and yellow modules.

**Tabel S4.** Hub gene expression scale in turquoise and yellow modules.

| Gene symbol | Log_FC_PCZ | LogFC_TFS | LogFC_PT |
| --- | --- | --- | --- |
| SUT2 | -1.44 | -1.698 | -1.406 |
| ARK2 | -1.389 | -1.655 | -1.493 |
| NTMC2T5.2 | -1.736 | -2.295 | -1.757 |
| ACA2 | -1.182 | -1.451 | -1.169 |
| HEMA2 | -1.678 | -2.17 | -1.651 |
| AT2G29320 | 1.217 | 1.262 | 1.619 |
| DOGT1 | 1.737 | 2.078 | 2.478 |
| ATH1 | 1.194 | 1.584 | 1.682 |

